# Toxicity Management in CAR T cell therapy for B-ALL: Mathematical modelling as a new avenue for improvement

**DOI:** 10.1101/049908

**Authors:** Shalla Hanson, David Robert Grimes, Jake P. Taylor-King, Benedikt Bauer, Pravnam I. Warman, Ziv Frankenstein, Artem Kaznatcheev, Michael J. Bonassar, Vincent L. Cannataro, Zeinab Y. Motawe, Ernesto A. B. F. Lima, Sungjune Kim, Marco L. Davila, Arturo Araujo

## Abstract

Advances in genetic engineering have made it possible to reprogram individual immune cells to express receptors that recognise markers on tumour cell surfaces. The process of re-engineering T cell lymphocytes to express *Chimeric Antigen Receptors* (CARs), and then re-infusing the CAR-modified T cells into patients to treat various cancers is referred to as CAR T cell therapy. This therapy is being explored in clinical trials - most prominently for B Cell Acute Lymphoblastic Leukaemia (B-ALL), a common B cell malignancy, for which CAR T cell therapy has led to remission in up to 90% of patients. Despite this extraordinary response rate, however, potentially fatal inflammatory side effects occur in up to 10% of patients who have positive responses. Further, approximately 50% of patients who initially respond to the therapy relapse. Significant improvement is thus necessary before the therapy can be made widely available for use in the clinic.

To inform future development, we develop a mathematical model to explore interactions between CAR T cells, inflammatory toxicity, and individual patients’ tumour burdens *in silico*. This paper outlines the underlying system of coupled ordinary differential equations designed based on well-known immunological principles and widely accepted views on the mechanism of toxicity development in CAR T cell therapy for B-ALL - and reports *in silico* outcomes in relationship to standard and recently conjectured predictors of toxicity in a heterogeneous, randomly generated patient population. Our initial results and analyses are consistent with and connect immunological mechanisms to the clinically observed, counterintuitive hypothesis that initial tumour burden is a stronger predictor of toxicity than is the dose of CAR T cells administered to patients.

We outline how the mechanism of action in CAR T cell therapy can give rise to such non-standard trends in toxicity development, and demonstrate the utility of mathematical modelling in understanding the relationship between predictors of toxicity, mechanism of action, and patient outcomes.

## I. INTRODUCTION

Traditionally three major modalities have have predominated in cancer therapy: surgery, radiation, and chemotherapy. While these interventions have lengthened survival times for cancer patients, they continue to undergo significant evolution in hopes of eventually finding a cure. Recently, efforts to find effective treatments without the severe side effects of traditional chemo-and radiation therapies have given rise to a new class of *targeted* drugs, which can distinguish potentially harmful cells from those that are likely healthy [1]. However, even with these continual advancements, cancer is still a challenge to treat. This is due in part to the fact that cancers often become resistant to therapies, and in part to the fact that even the best therapies often cause severe side effects that prevent high-dose or prolonged administration. These difficulties prompted interest in harnessing the power of the human immune system, which remarkably has both the ability to identify and attack specifically pathogenic cells and the ability to adapt to changing populations of cancer cells, to enhance treatments currently available for cancer patients. Today, *immunotherapy* presents an exciting new frontier in cancer therapy, replete with the possibility of reprogramming the human immune system itself to recognise and fight against cancer [2–4].

### Toxicity in cancer therapy

Chemo-and radiation therapies work by exploiting the fact that certain behaviours, in particular cell division, are more common in cancerous tissue than in healthy tissue. These therapies damage cells much more severely when they begin certain cellular processes, so if cancer cells undergo these processes more frequently than healthy cells do, more cancer cells are damaged by the therapy than healthy cells. Unfortunately, most healthy cells, to some extent, still undergo the same cellular processes exploited by traditional therapies and are thus indiscriminately damaged when they do.

Targeted agents, on the other hand, typically have a type of biochemical ‘switch’ which is turned ‘on’ when they encounter potentially malignant cells but remains ‘off’ otherwise. Ideally, the targeted agent is then toxic only to cells identified as harmful. The efficacy as well as the amount of toxicity caused by direct injury of healthy tissue thus depends on the specificity of the targeted agent. That is, the more malignant cells the agent is able to correctly identify as harmful and kill, the more effective the therapy. Conversely, the more healthy cells the agent is able to correctly identify as innocuous and leave alone, the lower the toxicity. Advances in targeted therapies have focused for some time on increasing the specificity of targeted agents, and include, for example, the addition of multiple biochemical switches to help identify malignant cells more accurately [5].

As promising as these therapies may be, this has unfortunately unveiled a second mechanism of toxicity development due, not to direct killing of healthy cells by the drug, but instead due to cancer cells being killed *too rapidly*. While killing cancer cells is precisely the intended outcome of these therapies, an extremely high rate of cell death within the body, regardless of cell type, can have severe consequences causing, for example seizures, renal failure, arrhythmias, and death [6]. This phenomenon is referred to as Tumour Lysis Syndrome (TLS) and has been increasing in incidence as cancer therapies become more and more targeted [7].

Another cell-death-associated side effect often seen in targeted immunotherapies is Cytokine Release Syndrome (CRS), which occurs when immune cells secrete massive quantities of inflammatory cytokines upon identifying pathogenic cells. Toxicities caused by CRS vary in severity but are often life-threatening, and include dangerously high fevers, precipitous drops in blood pressure, hypoxia, and neurological disorders [8].

Recent studies on toxicity associated with CD19-targeted CAR T cell therapy for B-ALL have reported very little correlation between toxicity and the dose of CAR T cells received by patients [9]. On the other hand, inflammatory toxicity *has* been reported to be much more highly correlated with individual patients’ tumour burdens than with the dose of CAR T cells administered [10, 11]. Nevertheless, despite the enigmatic toxicity profiles seen in these patients, a number of patients receiving the therapy have also shown sustained complete re-sponses with minimal toxicity development [12]. It is clear that in order for these types of outcomes to be possible for a broader patient population, we must first develop more a sophisticated understanding of toxicity development in targeted immunotherapies.

Nonetheless, due to the life-threatening results experienced by others, it is clear that major improvements must be made in toxicity management before such therapies can be safely administered. The following section reviews current research on mechanisms of action and toxicity development in CAR T cell therapy in B-ALL.

### CAR T cell therapy for B-ALL

The advent of stem-line engineering practices, which can endow cytotoxic (killer) T cells with the ability to identify and subsequently attack malignant cells [13], has propelled rapid progress in cancer treatment. This concept is the basis of CAR T cell therapy. Under this paradigm, T cells are collected from a patient or donor and genetically re-engineered to express Chimeric Antigen Receptors (CARs), which recognise tumour-specific markers called antigen. These cells are then stimulated to divide *in vitro* until a sufficiently large population (often on the order of 10^9^ cells) is produced, and are then intravenously infused back into the patient [9].

CAR T cell approaches for B cell tumours have been tentatively explored most often involve CARs specific for the B cell CD19 antigen [14–16]. CD19 is expressed on normal B cells, and is rarely lost due to mutations in B cell malignancies such as low-grade chronic lymphocytic leukemias (CLL), B cell non-Hodgkin’s lymphomas and more aggressive B-ALLs [17]. Various clinical trials have reported positive response rates of up to 90% [8, 18] in patients with B cell malignancies who received CAR T cell therapy. While this is encouraging, high relapse rates are still a significant problem. Follow-up studies on patients with relapsed diffuse large B cell lymphoma treated with anti-CD20 or anti-CD19 CAR T cells found that T cell persistence was no longer than seven days, likely related to a cellular anti-transgene immune response in some of the patients [19]. Though rare CD19-negative relapses have infrequently occurred in B-ALL patients whilst CAR T cells are still detectable[11], it is thought that the majority of B-ALL relapses are associated with disappearance of CAR T cells [20].

A deeper understanding of the dynamics leading to persistence or loss of CAR T cells is thus critical for further improvements in CAR T cell therapy. Unfortunately, the relapse rate among patients who re-ceive CAR T cell therapy is not the only problem. Roughly 10% of patients who respond to CAR T cell therapy develop a number of dangerous side effects, perhaps the most concerning of which is CRS [8, 21]. In the limited trials to date, CRS has been observed to be a common and potentially life-threatening side effect of CAR T cell therapy, this precludes approval for widespread use in the clinic.

To help overcome the limitations of CAR T cell therapy, circumvent relapse, and maximise treatment efficacy, mathematical modelling may be used to explore treatment dynamics *in silico*. Mathematical models can be developed to simulate different treatment regimes, and analysis of the coupled dynamics of both tumour regression and inflammatory side-effects can potentially be exceptionally useful. In this work, we establish a coupled ordinary differential equation model to simulate the interplay of tumour cells, inflammation, and several phenotypes of CAR T cells, and investigate the effects of different protocols and patient characteristics on treatment outcome.

## II. METHODOLOGY

### Model derivation

The majority of T cells can be subdivided into two varieties: *Cytotoxic* T cells - or commonly Cytotoxic T Lymphocytes (CTLs) - which are capable of directly killing tumour cells, and *Helper* T cells, which have the ability to enhance the killing potential of other T cells [22]. Further, both Cytotoxic and Helper T cell populations are themselves heterogeneous in phenotype, and recent studies have suggested that the phenotype of CAR T cells infused may modulate patient responses to the treatment [23]. In this investigation we focus on CAR-modified T cells, and initially confine ourselves to two distinct phenotypes, namely memory type and effector type, and recall that memory cells may later differentiate into effector cells [24, 25]. T cells with memory type traits have high proliferative capacity and are long lived. Effector type cells Cytotoxic and Helper T cells, by contrast, are short lived but much more effective, respectively, at either cell killing or enhancing the cell killing ability of other T cells than are memory type cells [26]. This heterogeneity is important in distinguishing population density from cytotoxicity, and thus therapeutic efficacy.

Here we present a brief simulation exploring the effects of CAR T cell therapy for B-ALL. The total populations of CAR-modified Helper T cells of memory and effector phenotype are denoted respectively by H_m_ and H_e_. Similarly, the populations of CAR-modified Cytotoxic T cells of memory and effector phenotype are denoted by C_m_ and C_e_. The background (non-CAR and non-B cell) endogenous lymphocyte population is denoted by *𝓛*. The total lymphocyte population is thus

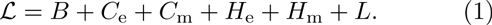

We will define the inflammatory response as *I*(*t*) and the B-ALL tumour burden as *B*(*t*). Initially and for simplicity we make the assumption that tumour growth is logistic. We note,however that for some patients, Gompertzian or other growth laws may give more accurate representations of tumour growth, and this will be considered in future work. For the present study, we write

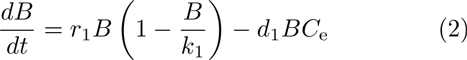

where *r*_1_ and *k*_1_ are constants of logistic growth, and *d*_1_ a constant associated with the killing potential of *C*_e_ cells.

The immune response is modelled to account for the fact that CAR-modified T cells secrete inflammatory molecules when they interact with CD19-positive B cells. While these inflammatory molecules decay naturally, if this decay is not fast enough, clinical interventions are sometimes necessary to mitigate inflammatory toxicity. Here we denote the immunosuppressant drug dose by *D*(*t*). We may now write the immune response to treatment and modulation as

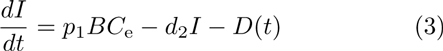

where *p*_1_, and *d*_2_ are production and decay constants respectively. Finally, we represent the respective populations of Cytotoxic and Helper T cells with the following system of ODEs:

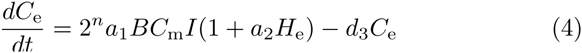

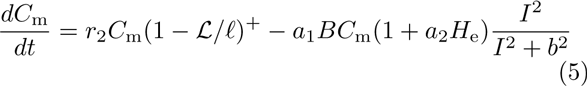

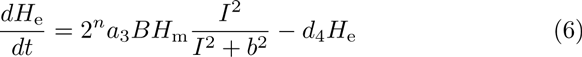

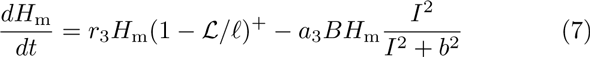

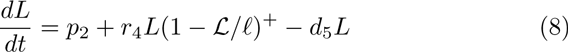

where *n* is the average number of divisions a differentiating T cell undergoes before becoming exhausted, *b* is the concentration of inflammatory cytokines which induces half-maximal differentiation rates in memory T cells, ℓ represents the normal concentration of lymphocytes, and *a*_i_, *p*_i_, *r*_i_, and *d*_i_ are rate constants associated respectively with activation, production, replication, and decay of the lymphocyte populations. The 2^n^ factor in the activation of memory T cells is used to account for the fact that T cells undergo successive divisions while differentiating from the memory to effector phenotype. The symbol + indicates the positive component, so *x*^+^ = max{*x*,0}. The term (1 – *𝓛/ℓ*)^+^ thus quantifies the degree (or absence) of lymphopenia. The Hill function, 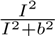, was chosen, firstly, to maintain a realistic bound on the activation rate of T cells, and secondly to include the presence of a homeostatic buffer which allows small changes of cytokine levels within the normal range without creating massive bursts in T cell activation.

Using these forms, we can simulate the effects of infusing different relative populations of CAR-modified T cells, immuno-suppression, and the interplay of different phenotypes in CAR T cell therapy-related toxicity. A simple outline of the model is shown in Figure 2.

**Figure 2:**
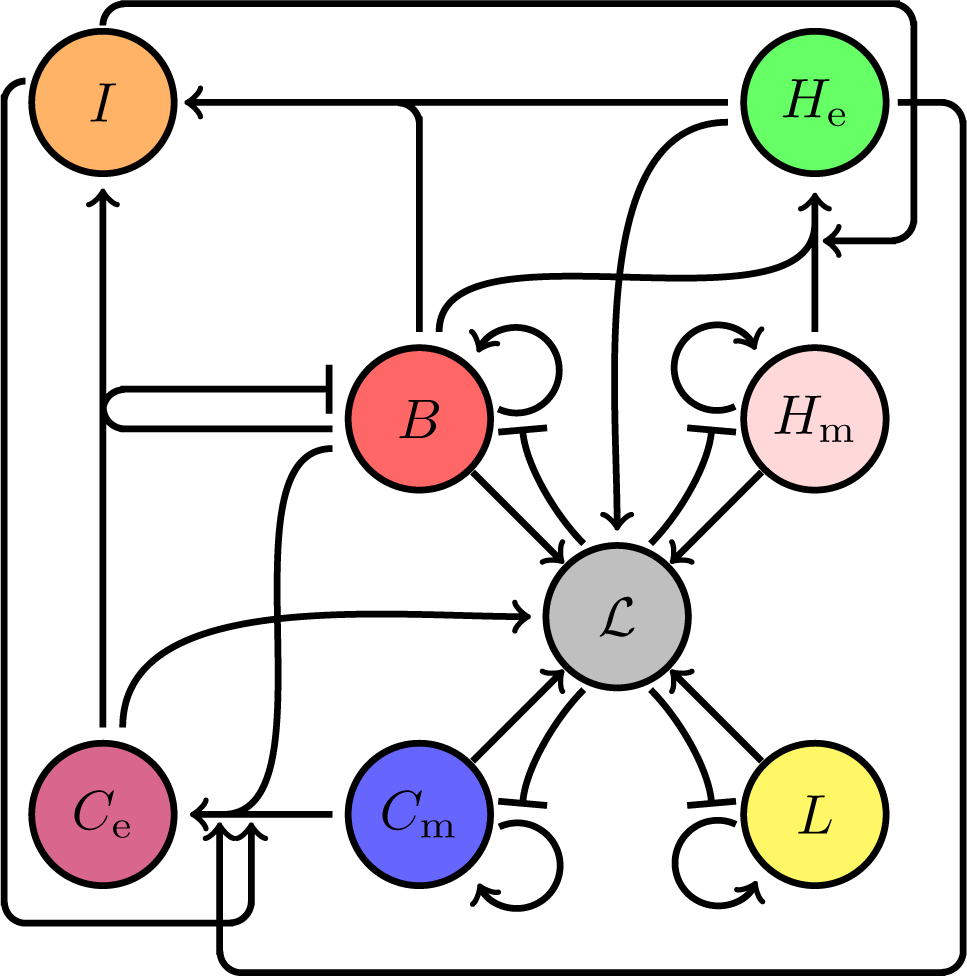
Schematic representation of key interactions. CAR-modified effector CTLs (C_e_), upon encountering B cells (B), secrete inflammatory cytokines (I) and cause lysis in B cells. Inflammation increases the rate at which CAR-modified memory CTLs (C_m_) and memory Helper T cells (H_m_) become effector cells. CAR-modified effector Helper T cells (H_e_) further increase the rate at which memory CTLs become effectors, and secrete inflammatory cytokines when they encounter B cells. The total number of lymphocytes (*𝓛*) increases when any one of the lymphocyte populations increases. Memory cells (H_m_, C_m_) and other endogenous lymphocytes (*𝓛*) undergo homeostatic division in response to lymphopenia, and B cells proliferate whenever L is below the carrying capacity.

### Simulation outline

The system of equations outlined in (1)–(8) was solved using an explicit 4^th^ order Runge– Kutta scheme. For the simulation in Figure 3, the initial conditions were chosen as listed in Table I, and parameters chosen as listed in Table II.

**Figure 3:**
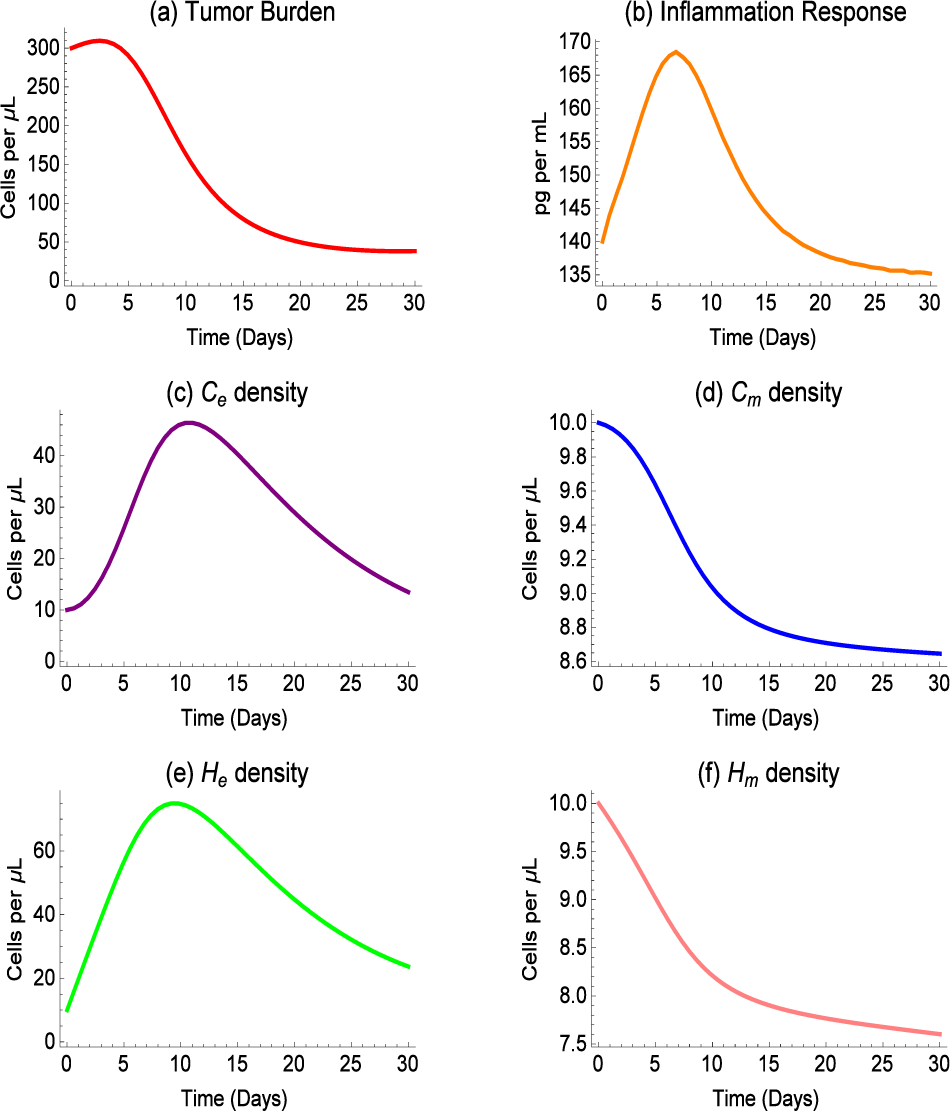
Simulated patient response. A simulated standard patient is administered CAR T cells on day 0 and monitored over 30 days. The tumour burden is depicted in (a), the inflammatory response in (b), and respective T cell populations are depicted in (c)-(e) inclusive. Simulation details are outlined in the text.

**Table I:** Initial Conditions

| Variable | Symbol |  |
| --- | --- | --- |
| Effector | $E(t)$ | |
| Target | $T(t)$ | 140 $\frac{pg}{mL}$ |
| CAR Memory CTLs | $C_m(t)$ | 10 $\frac{cells}{\mu L}$ |
| CAR Effector CTLs | $C_e(t)$ | 10 $\frac{cells}{\mu L}$ |
| CAR Memory Helper T cells | $H_m(t)$ | 10 $\frac{cells}{\mu L}$ |
| CAR Effector Helper T cells | $H_e(t)$ | 10 $\frac{cells}{\mu L}$ |
| Other lymphocytes | $L(t)$ | 1600 $\frac{cells}{\mu L}$ |

**Table II.**
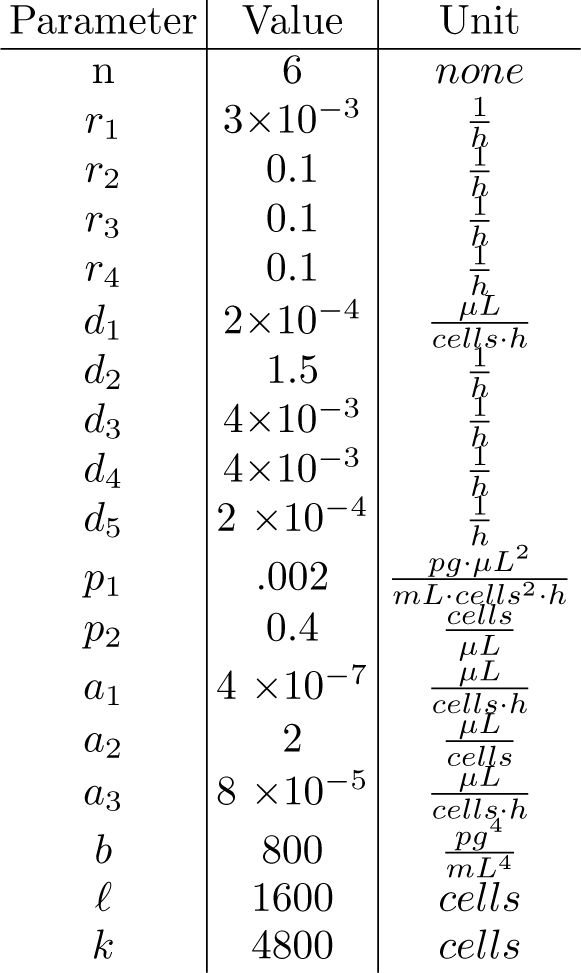
Parameter Values.

For clinical data simulations, a heterogeneous patient population (N = 200) was generated by randomly selecting parameter values and initial background endogenous lymphocyte concentrations either uniformly distributed between 75% and 125% of the mean values given above, or normally distributed about the means above, with standard deviations equal to 25% of the mean. Any non-physical (negative) parameter values generated were corrected to be within physical ranges. Initial tumour burden per patient was randomly selected from a uniform distribution between 1 and 1000 cells/μ*L*, and initial doses of each of the four CAR-modified T cell populations were similarly randomly selected from uniform distributions between 1 and 30 cells/μ*L* (such that the total CAR T cell dose administered per patient varies in the range of 1 to 120 cells/μ*L*).

## III. IN SILICO EXPERIMENTAL RESULTS

Detailed results for a single simulated patient are shown in Figure 3. For this patient, we see that CAR T cells undergo significant expansion and effector differentiation after infusion into patients with CD19^+^ BALL, and inflammatory responses peak between 5-10 days post-infusion concurrent with a rapid decrease in tumour burden. Subsequently, as the tumour burden declines, effector T cell populations decay and inflammation subsides. Further, we see that while the tumour burden may remain low for some time, the CAR T cell population continues to decline, predisposing the patient to relapse. These *in silico* results are consistent with clinical outcomes reported in [8, 10, 12].
Patient outcomes were also observed *in silico* in a randomly generated generated population (N=200) as outlined in the methodology. Consistent with the clinical results reported in [8–10, 12], we find that while inflammatory toxicity is only minimally correlated with the CAR T cell dose administered, it is in fact highly correlated with patients’ initial tumour burden (See Figure 4).

**Figure 4:**
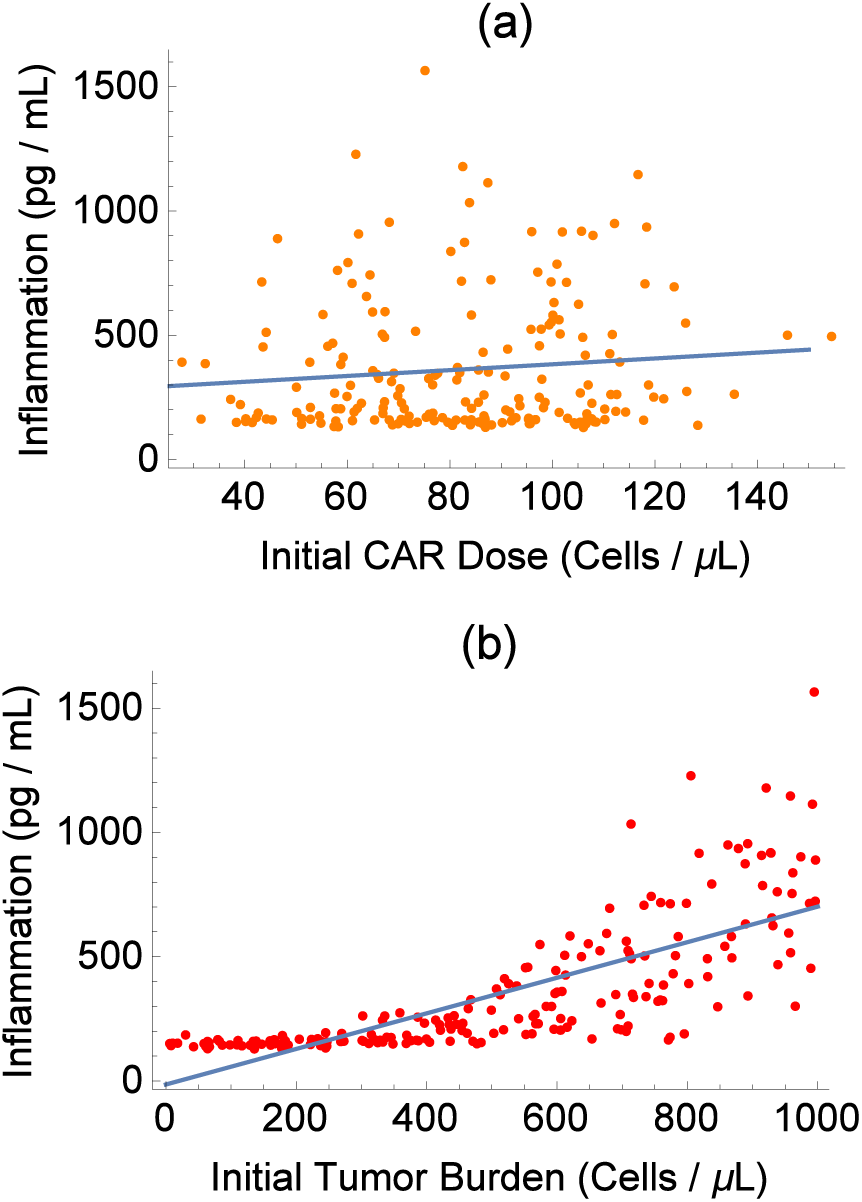
Simulated Clinical Data. Inflammatory molecules are secreted by CAR T cells when they interact with tumour cells, potentially resulting in life-threatening toxicities. (a) Maximum degree of inflammation experienced by patients (N=200) with initial CAR T cell dose in a simulated patient population. Linear best-fit is shown in blue, with a p-value 0.1412 suggesting no correlation between CAR load and toxicity. (b) Maximum degree of inflammation experienced by patients (N=200) with initial tumour burden in a simulated patient population. Linear best-fit is shown in blue, with a p-value 5.3 ×10^−38^, indicating extremely high correlation between tumour burden and toxicity. Data shown above was simulated from a patient population with individual characteristics (parameter values) chosen to be uniformly distributed about the mean. Simulations from a normally distributed patient population were also carried out and yield similar results.

## IV. DISCUSSION AND CONCLUSION

The observation that efficacy and toxicity are more highly correlated with tumour burden than with the initial dose of CAR T cells administered to patients [8– 10, 12] is contrary to what is observed with most pharmaceuticals [27]. Specifically, in non-targeted therapies, including chemo-and radiotherapy, toxicity results primarily from direct injury of healthy cells (or their daughter cells) by the therapeutic agent alone. As a result, the toxicity experienced by patients is independent of the size or density of the tumour. On the other hand, in CAR T cell therapy for B-ALL, toxicity results primarily from indirect injury of healthy tissue due to excessive cytokine secretion by CAR T cells when they encounter tumour cells (as depicted in Figure 1).

**Figure 1:**
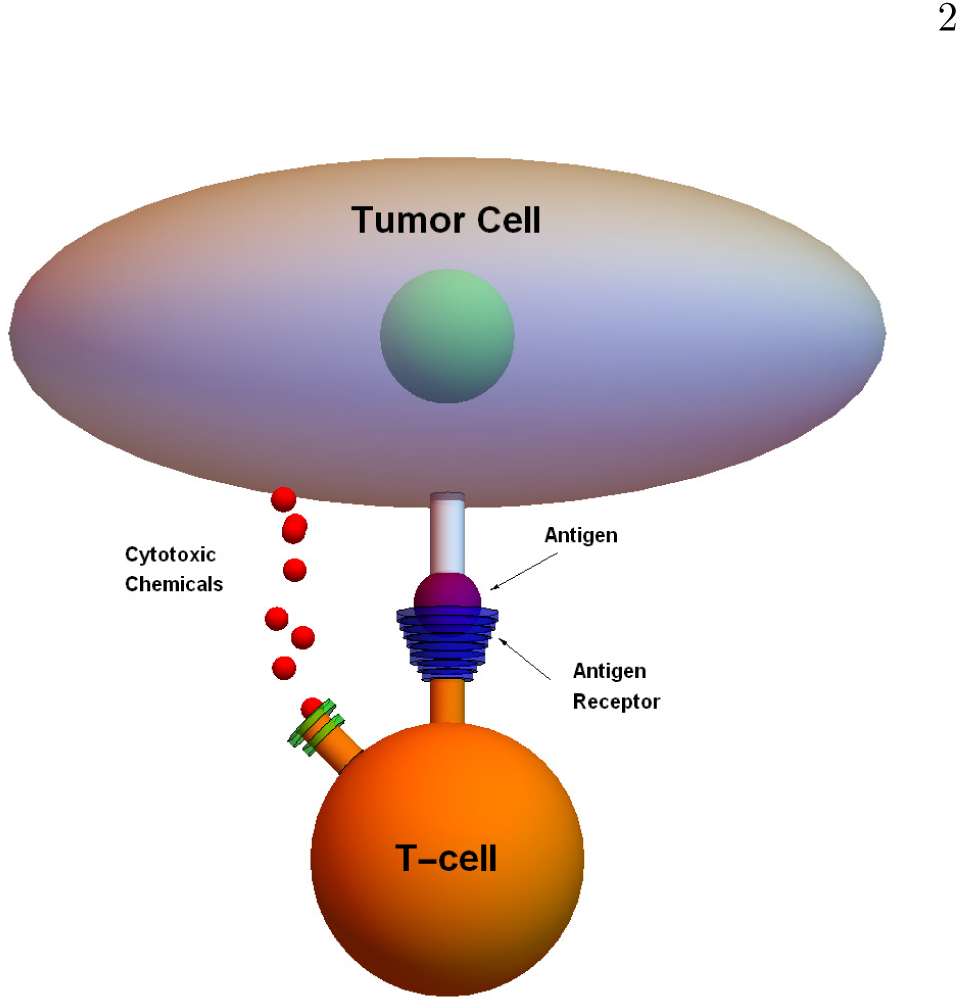
Chimeric Antigen Receptor (CAR)-modified T cell interacting with its cognate antigen on the surface of a tumour cell. A CAR-modified Cytotoxic T cell re-engineered to express a receptor specific to a tumour cell antigen identifies a tumour cell and secretes cytotoxic molecules which cause lysis of the tumour cell.

Our results demonstrate that this well-known mechanism of inflammatory cytokine secretion [28] is sufficient to give rise to tumour-burden-correlated toxicity as suggested in [8]. Our results also support the hypotheses of [9] that CAR T cell dose alone correlates poorly with the degree of toxicity experienced by patients, and along with the work of [8–10, 12], suggests that to best take advantage of this exciting new technology, we must also take steps to advance our practices for managing toxicity in targeted immunotherapies.

Mathematical analysis is a valuable tool for deepening our understanding of how the mechanisms of action of particular therapies connects to patient outcomes in the clinic [29]. It can potentially offer insight into how protocols may be modified based on individual patient characteristics to decrease toxicity, increase persistence of CAR T cell populations, and potentially decrease relapse rates in B-ALL. While the simple model presented here is at a very early stage of development, it illustrates the utility of mathematical modelling. Future work will focus on refining this approach, increasing precision in model predictions through experimental validation, and expansion to solid and other more heterogeneous tumours for which a single targetable antigen is not preserved in the tumour cell population.

## V. ACKNOWLEDGEMENTS

The group would like to kindly thank H. Lee Moffitt Cancer Center and Research Institute and in particular Alexander R. A. Anderson for the organisation and funding of the 5^th^ IMO workshop: Immune Cancer; and also Jean Clairambault and David Basanta for generously offering their guidance and expertise. SH received funding in part from the NSF under grant reference number DMS-0943760; and in part from NESCent, TriCEM, the Embassy of France in the United States, and INRIA-Paris. JPT-K received funding from the EPSRC under grant reference number EP/G037280/1. DRG is funded by CR:UK grants C5255/A15935 and C53469/A19834. B.B is funded by the Max Planck Society. EABFLis funded by the NSF under Grant DMS-1115865. AA is funded by DoD grant 14091003.

## References

[1] Devita VT, Chu E. AACR Centennial Series A History of Cancer Chemotherapy. Cancer Research. 2008;68(21):8643–8653.

[2] Currie G. Eighty Years of Immunotherapy/A Review of Immunological Methods Used for the Treatment of Human Cancer. British Journal of Cancer. 1972;26(3):141–153.

[3] Rosenberg Sa, Spiess P, Lafreniere R. A new approach to the adoptive immunotherapy of cancer with tumor-infiltrating lymphocytes. Science (New York, NY). 1986;233(4770):1318–1321.

[4] Wolchok JD, Hoos A, Day SO, Weber JS, Hamid O, Lebbé C, et al. Guidelines for the Evaluation of Immune Therapy Activity in Solid Tumors: Immune-Related Response Criteria. Clinical Cancer Research. 2009;15(23):7412–7420.

[5] Sun LL, Ellerman D, Mathieu M, Hristopoulos M, Chen X, Li Y, et al. Anti-CD20 / CD3 T cell dependent bispecific antibody for the treatment of B cell malignancies. Immunotherapy. 2015;7(287):1–10′.

[6] Howard S, Jones D, Pui C. The Tumor Lysis Syndrome. The New England Journal of ‥‥ 2011;364(19):1844–1854. Available from: http://www.nejm.org/doi/full/10.1056/nejmra0904569.

[7] McBride A, Westervelt P. Recognizing and managing the expanded risk of tumor lysis syndrome in hematologic and solid malignancies. Journal of hematology & oncology. 2012;5(1):75. Available from: http://www.pubmedcentral.nih.gov/articlerender.fcgi?artid=3544586{&}tool=pmcentrez{&}rendertype=abstract.

[8] Davila ML, Riviere I, Wang X, Bartido S, Park J, Curran K, et al. Efficacy and toxicity management of 19-28z CAR T cell therapy in B cell acute lymphoblastic leukemia. Science translational medicine. 2014;6(224):224ra25. Available from: http://www.pubmedcentral.nih.gov/articlerender.fcgi?artid=4684949{&}tool=pmcentrez{&}rendertype=abstract.

[9] Davila ML, Brentjens R, Wang X, Rivière I, Sadelain M. How do CARs work?: Early insights from recent clinical studies targeting CD19. Oncoimmunology. 2012;1(9):1577–1583. Available from: http://www.pubmedcentral.nih.gov/articlerender.fcgi?artid=3525612{&}tool=pmcentrez{&}rendertype=abstract.

[10] Brentjens RJ, Davila ML, Riviere I, Park J, Wang X, Cowell LG, et al. CD19-targeted T cells rapidly induce molecular remissions in adults with chemotherapy-refractory acute lymphoblastic leukemia. Science translational medicine. 2013;5(177):177ra38. Available from: http://www.pubmedcentral.nih.gov/articlerender.fcgi?artid=3742551{&}tool=pmcentrez{&}rendertype=abstract.

[11] Lee DW, Kochenderfer JN, Stetler-Stevenson M, Cui YK, Delbrook C, Feldman SA, et al. T cells expressing CD19 chimeric antigen receptors for acute lymphoblastic leukaemia in children and young adults: A phase 1 dose-escalation trial. The Lancet. 2015;385(9967):517–528. Available from: http://dx.doi.org/10.1016/S0140-6736(14)61403-3.

[12] Brudno JN, Somerville RPT, Shi V, Rose JJ, Halverson DC, Fowler DH, et al. Allogeneic T Cells That Express an Anti-CD19 Chimeric Antigen Receptor Induce Remissions of B-Cell Malignancies That Progress After Allogeneic Hematopoietic Stem-Cell Transplantation Without Causing Graft-Versus-Host Disease. Journal of Clinical Oncology. 2016;Available from: http://jco.ascopubs.org/cgi/doi/10.1200/JCO.2015.64.5929.

[13] Sadelain M, Brentjens R, Rivière I. The promise and potential pitfalls of chimeric antigen receptors. Current Opinion in Immunology. 2009;21(2):215–223.

[14] Porter DL, Levine BL, Kalos M, Bagg A, June CH. Chimeric Antigen ReceptorModified T Cells in Chronic Lymphoid Leukemia. New England Journal of Medicine. 2011;365(8):725–733.

[15] Kalos M, Levine BL, Porter DL, Katz S, Grupp SA, Bagg A, et al. T Cells with Chimeric Antigen Receptors Have Potent Antitumor Effects and Can Establish Memory in Patients with Advanced Leukemia. LEUKEMIA. 2011;3(95):95ra73—95ra73.

[16] Grupp SA, Kalos M, Barrett D, Aplenc R, Porter DL, Rheingold SR, et al. Chimeric Antigen Recep-torModified T Cells for Acute Lymphoid Leukemia. 2013;368(16):1509–1518.

[17] Wang K, Wei G, Liu D. CD19: a biomarker for B cell development, lymphoma diagnosis and therapy. Experimental hematology & oncology. 2012;1(1):36. Available from: http://ehoonline.biomedcentral.com/articles/10.1186/2162-3619-1-36.

[18] Maude SL, Barrett D, Teachey DT, Grupp SA. Managing cytokine release syndrome associated with novel T cell-engaging therapies. Cancer journal (Sudbury, Mass). 2014;20(2):119.

[19] Jensen MC, Popplewell L, Cooper LJ, DiGiusto D, Kalos M, Ostberg JR, et al. Antitransgene rejection responses contribute to attenuated persistence of adoptively transferred CD20/CD19-specific chimeric antigen receptor redirected T cells in humans. Biology of Blood and Marrow Transplantation. 2010;16(9):1245–1256.

[20] Dai H, Wang Y, Lu X, Han W. Chimeric Antigen Receptors Modified T-Cells for Cancer Therapy. Journal of the National Cancer Institute. 2016;108(7):1–15.

[21] Kochenderfer JN, Dudley ME, Feldman SA, Wilson WH, Spaner DE, Maric I, et al. B-cell depletion and remissions of malignancy along with cytokine-associated toxicity in a clinical trial of anti-CD19 chimeric-antigen-receptor transduced T cells. Blood. 2011;119(12):2709– 2720.

[22] Broere F, Apasov SG, Sitkovsky MV, Eden WV. Principles of Immunopharmacology. Principles of Immunopharmacology. 2011;p. 15–28. Available from: http://link.springer.com/10.1007/978-3-0346-0136-8.

[23] Ding Zc, Huang L, Blazar BR, Yagita H, Mellor AL, Munn DH, et al. Polyfunctional CD4 + T cells are essential for eradicating advanced B-cell lymphoma after chemotherapy Polyfunctional CD4 T cells are essential for eradicating advanced B-cell lymphoma after chemotherapy. Blood. 2012;120(11):2229–2239.

[24] Waddington CH. The Strategy of the Genes: A Discussion of Some Aspects of Theoretical Biology. London: George Allen & Unwin; 1957.

[25] Graef P, Buchholz VR, Stemberger C, Flossdorf M, Henkel L, Schiemann M, et al. Serial Transfer of Single-Cell-Derived Immunocompetence Reveals Stemness of CD8+ Central Memory T Cells. Immunity. 2014;41:116– 126.

[26] Kaech SM, Cui W. Transcriptional control of effector and memory CD8+ T cell differentiation. Nat Rev Immunol. 2012;12(11):749–761. Available from: http://dx.doi.org/10.1038/nri3307.

[27] Waddell WJ. History of dose response. The Journal of Toxicological Sciences. 2010;35(1):1–8.

[28] Varadarajan N, Julg B, Yamanaka YJ, Chen H, Ogunniyi aO, McAndrew E, et al. A high-throughput single-cell analysis of human CD8 + T cell functions reveals discordance for cytokine secretion and cytolysis. Journal of Clinical Investigation. 2011;121(11):4322– 4331. Available from: http://www.scopus.com/inward/record.url?eid=2-s2.0-80555148949{&}partnerID=40{&}md5=94912cd57ca14270f5e27205ebd00e13.

[29] Anderson ARa, Quaranta V. Integrative mathematical oncology. Nature reviews Cancer. 2008;8(3):227–234.

